# Repertoire of naturally acquired maternal antibodies transferred to infants for protection against shigellosis

**DOI:** 10.1101/2021.05.21.445178

**Authors:** Esther Ndungo, Liana R. Andronescu, Andrea G Buchwald, Jose M. Lemme-Dumit, Patricia Mawindo, Neeraj Kapoor, Jeff Fairman, Miriam K. Laufer, Marcela F. Pasetti

## Abstract

*Shigella* is the second leading cause of diarrheal diseases, accounting for >200,000 infections and >50,000 deaths in children under 5 years of age annually worldwide. The incidence of *Shigella*-induced diarrhea is relatively low during the first year of life and increases substantially, reaching its peak between 11 to 24 months of age. This epidemiological trend hints at an early protective immunity of maternal origin and an increase in disease incidence when maternally acquired immunity wanes. The magnitude, type, antigenic diversity, and antimicrobial activity of maternal antibodies transferred via placenta that can prevent shigellosis during early infancy are not known. To address this knowledge gap, *Shigella-*specific antibodies directed against the lipopolysaccharide (LPS) and virulence factors (IpaB, IpaC, IpaD, IpaH, and VirG), and antibody-mediated serum bactericidal (SBA) and opsonophagocytic killing antibody (OPKA) activity were measured in maternal and cord blood sera from a longitudinal cohort of mother-infant pairs living in rural Malawi. Protein-specific (very high levels) and *Shigella* LPS IgG were detected in maternal and cord blood sera; efficiency of placental transfer was 100% and 60%, respectively, and had preferential IgG subclass distribution (protein-specific IgG1 > LPS-specific IgG2). In contrast, SBA and OPKA activity in cord blood was substantially lower as compared to maternal serum and varied among *Shigella* serotypes. LPS was identified as the primary target of SBA and OPKA activity. Maternal sera had remarkably elevated *Shigella flexneri* 2a LPS IgM, indicative of recent exposure. Our study revealed a broad repertoire of maternally acquired antibodies in infants living in a *Shigella*-endemic region and highlights the abundance of protein-specific antibodies and their likely contribution to disease prevention during the first months of life. These results contribute new knowledge on maternal infant immunity and target antigens that can inform the development of vaccines or therapeutics that can extend protection after maternally transferred immunity wanes.

## Introduction

*Shigella* spp. are major contributors of the global diarrheal disease burden, accounting for more than 250 million cases and 200,000 deaths annually (1, 2). The most affected are children under 5 years of age living in low- and middle-income countries (LMIC) (2, 3). Though usually self-limiting, repeated bouts of disease result in debilitating sequalae including malnutrition, stunted growth, and deficits in immune and cognitive development (3, 4). The preeminence of multidrug resistant *Shigella* strains globally makes the development of vaccines and therapeutics a compelling priority (5). Because the burden of disease disproportionately affects young children, a clear understanding of the elements and immune mechanisms that can protect this group is necessary to inform the development of efficacious vaccines or prophylaxes.

Most of what is known about *Shigella* immunity has been learned from infections in adults. Individuals living in endemic regions acquire natural immunity from repeated exposure (6-9). While there is no definitive immune correlate of protection against shigellosis, serum IgG against the *Shigella* surface-exposed lipopolysaccharide (LPS) has been associated with reduced risk of infection with serotype-matching strains in early field trials [reviewed in Ref (10)]. We have presented evidence that serum IgG specific for the *Shigella* invasion plasmid antigen (Ipa) B and the virulence protein VirG (IcsA) were associated with reduced risk of infection in a controlled human infection model (CHIM) study (11). In the same experimentally infected adult volunteers, complement-dependent serum bactericidal (SBA) and opsonophagocytic killing (OPKA) activity were identified as functional attributes of *Shigella*-specific antibodies associated with clinical protection (11).

Children living in endemic regions produce serum LPS- and Ipa-specific IgG in response to *Shigella* infection, and the magnitude of these responses increases progressively through adulthood (6, 9, 12, 13). Multiple surveillance studies have reported consistently that the rate of *Shigella* infection is relatively low during the first months of life, but gradually increases and reaches its peak during the second year of life (14, 15). The shielding of infants from *Shigella*-induced diarrhea early in life (although they may still suffer from other enteric infections such as rotavirus), hints at a putative pathogen-specific protection afforded by maternal immunity, i.e. antibodies transferred via placenta and the immune components of breast milk (16). Studies of transplacental antibody transfer against other pathogens have shown that this process—termed placental “sieving” (17)—is regulated and selective, antigen-dependent (18, 19), and favors transfer of antibodies with specific biophysical features that make them most effective in the immature neonatal immune system (17). Information on antigen-specificity, magnitude, subclass distribution, and function of *Shigella* antibodies in mothers and infants and the process of placental transfer has been lacking. Here, we characterized the specificity and antimicrobial function of *Shigella*-specific antibodies in mothers and their infants at birth in a longitudinal cohort from rural Malawi. The magnitude of serum IgG (and IgG subclasses) specific for LPS against two *Shigella* serotypes, *S. flexneri* 2a and *S. sonnei*, and conserved *Shigella* proteins (virulence factors) IpaB, IpaC, IpaD, IpaH, and VirG were determined. SBA and OPKA levels and the target antigen mediating complement- and phagocytic-effector functions were investigated. Finally, correlative analysis and comparison with protective thresholds were conducted to identify unique features and the potential in vivo antimicrobial activity of *Shigella* antibodies in mother-infant pairs.

## Results

### Cohort characteristics

This study utilized a mother-infant cohort from a malaria surveillance study in Malawi. Participants were enrolled from rural villages in Chikwawa in the southern region. Out of 108 mother and infant pairs enrolled, 63 pairs were analyzable (Figure 1). Cord blood was collected at birth during vaginal deliveries at the Mfera Clinic; mothers requiring caesarian deliveries were referred to a hospital. Maternal blood was obtained at recruitment within 3 months of delivery. Characteristics of the cohort are summarized in Table 1. Mean age of the mothers was 26.8 years (17-43 years). Median maternal parity was 3 (0-8). Among the infants, 36 (57%) were female and 5 (8%) had a low birth weight (less than 2.5kg). As for the season of birth, 21 (33%) were born during the rainy season (November – April).

**Table 1.** Baseline Characteristics of 63 Participating Mother-Infant Pairs.

| Characteristic | Value |
| --- | --- |
| Maternal characteristics |  |
| Age, y, median (range) | 26 (17-43) |
| Parity, median (range) | 3 (0-8) |
| Infant characteristics |  |
| Female sex (%) | 36 (57%) |
| Birth weight, kg, median (range) | 3.1 (1.5-4.3) |
| < 2.5 kg (low birth weight), n (%) | 5 (8%) |
| Twin births, n (%) | 3(4.8%) |
| Born during rainy season (November- April), n (%) | 21 (33%) |

**Figure 1.**
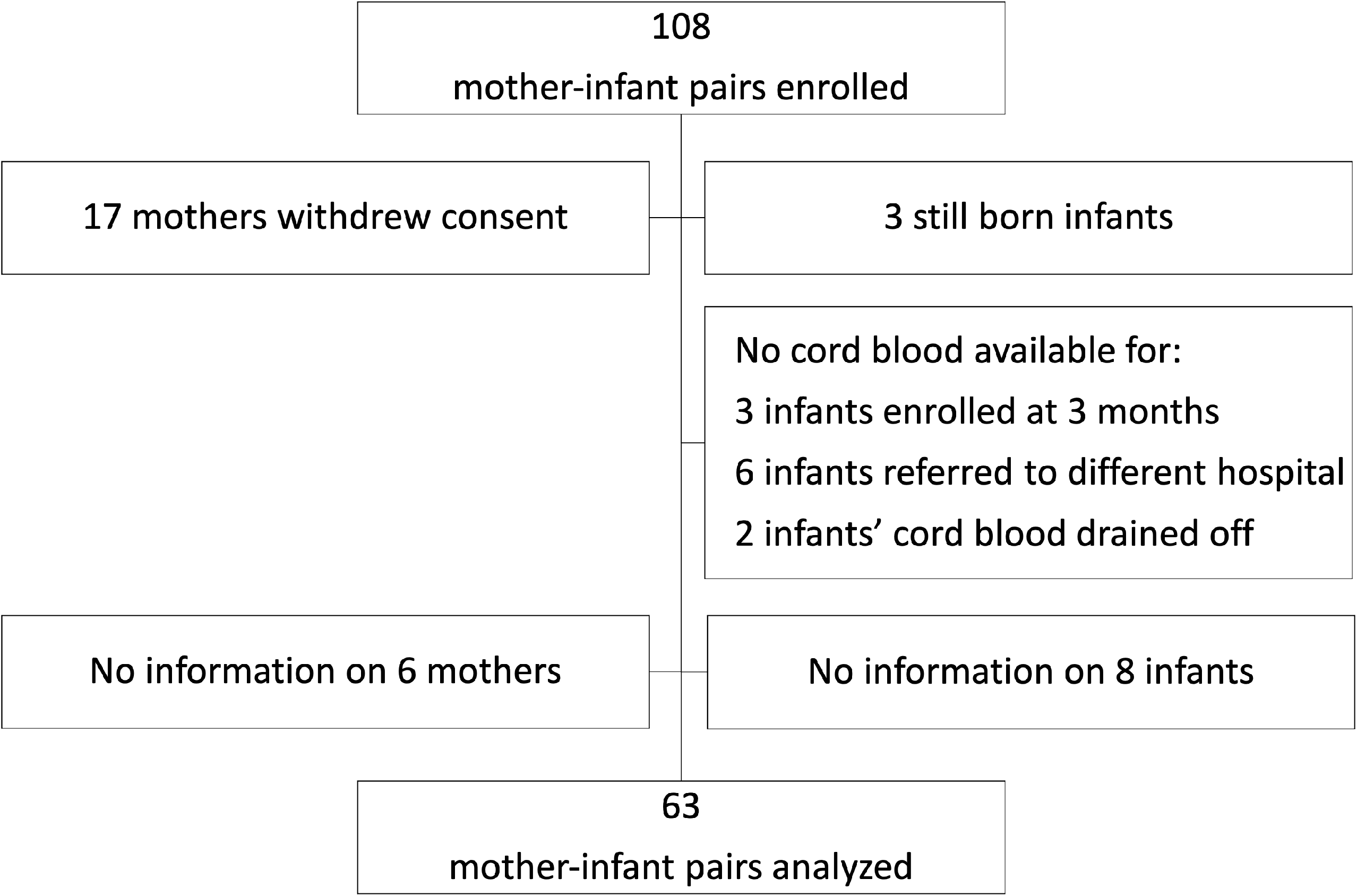
Flowchart showing selection of paired maternal-infant samples.

### *Shigella* antigen-specific IgG in mothers and their newborns

Naturally acquired *Shigella*-specific antibodies were determined in paired maternal and cord blood sera. The antigenic repertoire analysis focused on *S. flexneri* 2a and *S. sonnei*; these species had been attributed the highest incidence of moderate-to-severe diarrhea (MSD) in <5-year-old children (37.8% and 13.5%, respectively) in Malawi-neighboring Mozambique by the Global Enteric Multicenter Study (GEMS) (20); precise information on *Shigella* prevalence in Malawi is not available. *S. flexneri* 2a and *S. sonnei* LPS-specific IgG titers in maternal sera were significantly higher as compared to those in cord blood (Figure 2A). The median cord-blood to maternal *S. flexneri* 2a and *S. sonnei* LPS IgG transfer ratios were 0.51 and 0.57 respectively (Figure 2B, Table 2), indicating low transplacental sieving efficiency of LPS IgG.

**Table 2.** *Shigella* antigen-specific titers.

| Antigen | Cord blood GMT<br>EU/mL (range) | Maternal serum GMT<br>EU/mL (range) | Transfer ratio<br>Median (range) |
| --- | --- | --- | --- |
| *S. flexneri* 2a LPS | 1,752 (120-26,789) | 3,491 (153-123,844) | 0.51 (0.05-2.20) |
| *S. sonnei* LPS | 1,264 (45-19,002) | 2,109 (135-32,727) | 0.57 (0.23-3.38) |
| IpaB | 77,134 (7,274-419,898) | 73,737 (6,818-324,773) | 1.06 (0.27-2.00) |
| IpaC | 21,856 (279-125,268) | 21,732 (292-127,040) | 1.02 (0.34-2.02) |
| IpaD | 9,761 (84-92,804) | 10,653 (81-112,576) | 0.93 (0.26-2.08) |
| IpaH | 19,890 (1,224-122,206) | 21,156 (1,067-145,110) | 0.97 (0.21-1.98) |
| VirG | 14,840 (3,038-63,877) | 15,726 (2,003-75,273) | 0.96 (0.29-3.71) |

**Figure 2.**
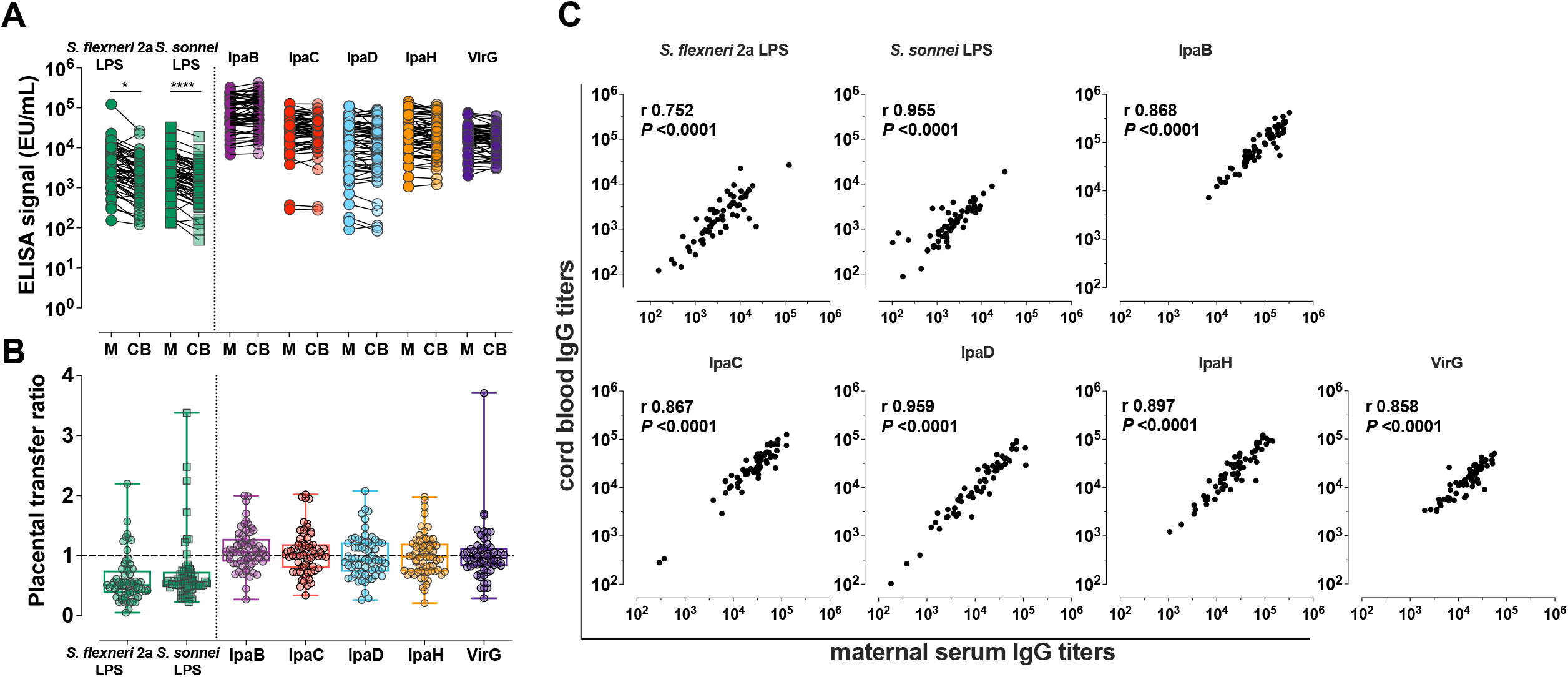
*Shigella*-specific maternal antibody repertoire and placental transfer efficiency. (A) IgG against *Shigella* LPS and protein antigens in maternal (M) and cord blood (CB) sera. Symbols represent individual titers. Asterisks indicate statistically significant differences determined by paired t-test; * *P* < 0.05, **** *P* < 0.0001. (B) Placental transfer ratios (cord blood titer/maternal titer) for antibodies against each antigen. Whiskers show minimum and maximum values. Line is at ratio = 1. (C) Associations between maternal and cord blood titers for each antigen. Pearson’s r and *P* values are shown within each graph.

Maternal and cord blood IgG levels against five different *Shigella* virulence factors: IpaB, IpaC, IpaD, IpaH, and VirG and their placental transfer efficiency were also determined (Figure 2A and 2B). High levels of circulating IgG specific for all protein antigens were detected in both maternal and cord blood sera, which far surpassed the levels of IgG against LPS (Figure 2A). Likewise, placental transfer of protein-specific maternal IgG was more efficient than the transfer of LPS-specific IgG; median transfer ratios were: 1.06, 1.02, 0.93, 0.97, and 0.96 for anti-IpaB, - IpaC, -IpaD, -IpaH, and -VirG antibodies, respectively (Figure 2B and Table 2). Despite differences in transfer efficiency between LPS- and protein-specific antibody titers, there was a significant and positive linear correlation between maternal and cord blood IgG levels for all antigens, confirming the selective and distinct regulation of placental antibody transport (Figure 2C). Geometric mean titers (GMT), median transfer ratios, and range of maternal and cord serum titers are summarized in Table 2.

We also examined whether maternal and infant variables collected in our study (Table 1) influenced maternal-infant *Shigella* antibody transfer. Maternal serum IgG titer was negatively associated with antibody transfer efficiency (Supplementary table 1). The same phenomenon has been observed in previous studies of maternal antibody transfer (18, 21, 22) and has been attributed to the saturation of placental Fc receptors at high maternal IgG concentrations which limits the amount of transferred antibodies (Reviewed in (23)). There was no significant correlation between the IgG transfer ratios and the maternal age, parity, gestational age, and infant birthweight (Supplementary table 1).

### *Shigella* protein- and LPS-specific IgG subclass placental transfer

Placental transport of maternal antibodies is primarily mediated through binding to the neonatal Fc receptor (FcRn) expressed in the syncytiotrophoblast (24, 25). Qualitative differences in the Fc structure, such as in the human IgG subclasses, can influence FcRn binding and placental transfer (26). We therefore explored IgG subclass distribution of *Shigella*-specific antibodies as a contributor to the observed differences in IgG transfer efficiency.

As with total IgG titers, the protein-specific IgG subclass repertoire had common features, which differed from those of IgG subclasses against LPS. In both the mothers and their infants, IgG1 was the most abundant subclass against all protein antigens, followed by IgG2 and IgG3 (Figure 3A and Supplementary table 2). In contrast, IgG2 was the most abundant subclass against both *S. flexneri* 2a and *S. sonnei* LPS. IgG4 titers were generally the lowest for all antigens tested (Figure 3A and Supplementary table 2). For protein antigens, IgG1 and IgG4 had the highest cord-blood:maternal median transfer ratios: 0.97-1.14 and 1.25-1.72, respectively (Figure 3B and Supplementary table 2).

**Figure 3.**
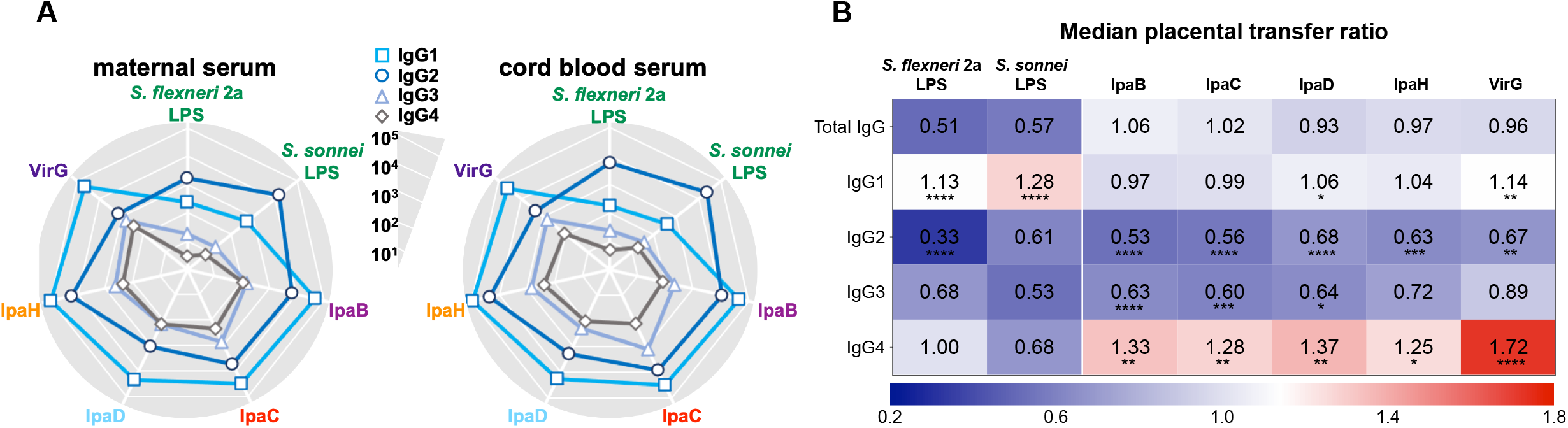
IgG subclass distribution of maternal and infant placentally-acquired antibodies. (A) Radar plots depict IgG1, IgG2, IgG3, and IgG4 levels against *Shigella* antigens measured in maternal and cord blood sera diluted at 1:100. Symbols represent the geometric mean Electro Chemiluminescence (ECL) Signal. (B) Heatmap representing median placental transfer ratios (cord blood titer/maternal titer) for *Shigella* antigen-specific IgG and IgG subclasses. Statistically significant differences between the transfer ratio of each IgG subclass compared to the total IgG transfer ratio were determined by one-way ANOVA with Dunnett’s post-test correction following ROUT analysis to exclude outliers; * *P* < 0.05, ** *P* < 0.01, *** *P* < 0.001, **** *P* < 0.0001.

For LPS antigens, IgG1 also exhibited the highest median transfer ratios: 1.13 and 1.28 for *S. flexneri* 2a and *S. sonnei*, respectively (Figure 3B and Supplementary table 2). Median transfer ratios for LPS IgG2 were lower, the lowest being for IgG2 against *S. flexneri* 2a. The lower transfer efficiency of LPS-specific IgG2 explains the modest levels of LPS IgG in the cord blood despite their abundance in maternal circulation. The median transfer ratios for antigen-specific IgG compared to IgG subclasses are represented in a heatmap (Figure 3B).

### Functional capacity of placentally transferred *Shigella*-specific antibodies

In addition to antibody specificity through direct antigen binding, we examined the functional capacity of maternal and placentally-acquired antibodies to render complement-dependent bactericidal and opsonophagocytic activity. SBA and OPKA activity were detected in both maternal and newborn sera. Maternal SBA and OPKA titers against *S. flexneri* 2a were significantly higher as compared to those against *S. sonnei* (GMT 26,944 compared to 306, respectively). Maternal SBA and OPKA titers specific for *S. flexneri* 2a were also significantly higher in maternal sera as compared to those of cord blood (Figure 4A and B); the median transfer ratios were 0.02 and 0.03, respectively (Figure 4A and B, and Table 3). In contrast, SBA and OPKA titers against *S. sonnei* in maternal and infant sera were comparable (Figure 4A and B); the median transfer ratios were 0.91 and 0.75 (Figure 4A and B and Table 3). It was noticed that while maternal and cord blood functional antibody titers against *S. sonnei* were strongly correlated, those against *S. flexneri* 2a were not (Figure 4C and D). The discrepancy in functional antibody titer against *S. flexneri* 2a between mothers and infants prompted us to investigate the specificity and type of antibodies involved in bactericidal and opsonophagocytic killing.

**Table 3.** *Shigella* functional antibody (SBA and OPKA) titers.

| Antigen | Cord blood<br>GMT (range) | Maternal serum<br>GMT (range) | Transfer ratio<br>Median (range) |
| --- | --- | --- | --- |
| *S. flexneri* 2a SBA | 1,227 (200-113,628) | 44,076 (6,889-1,278,483) | 0.02 (0.00-2.78) |
| *S. flexneri* 2a OPKA | 895 (100-178,424) | 26,944 (2,770-596,279) | 0.03 (0.00-1.77) |
| *S. sonnei* SBA | 832 (200-13,536) | 1,017 (200-17,916) | 0.91 (0.19-13.61) |
| *S. sonnei* OPKA | 245 (100-2,468) | 357 (100-2,460) | 0.75 (0.18-3.03) |

**Figure 4.**
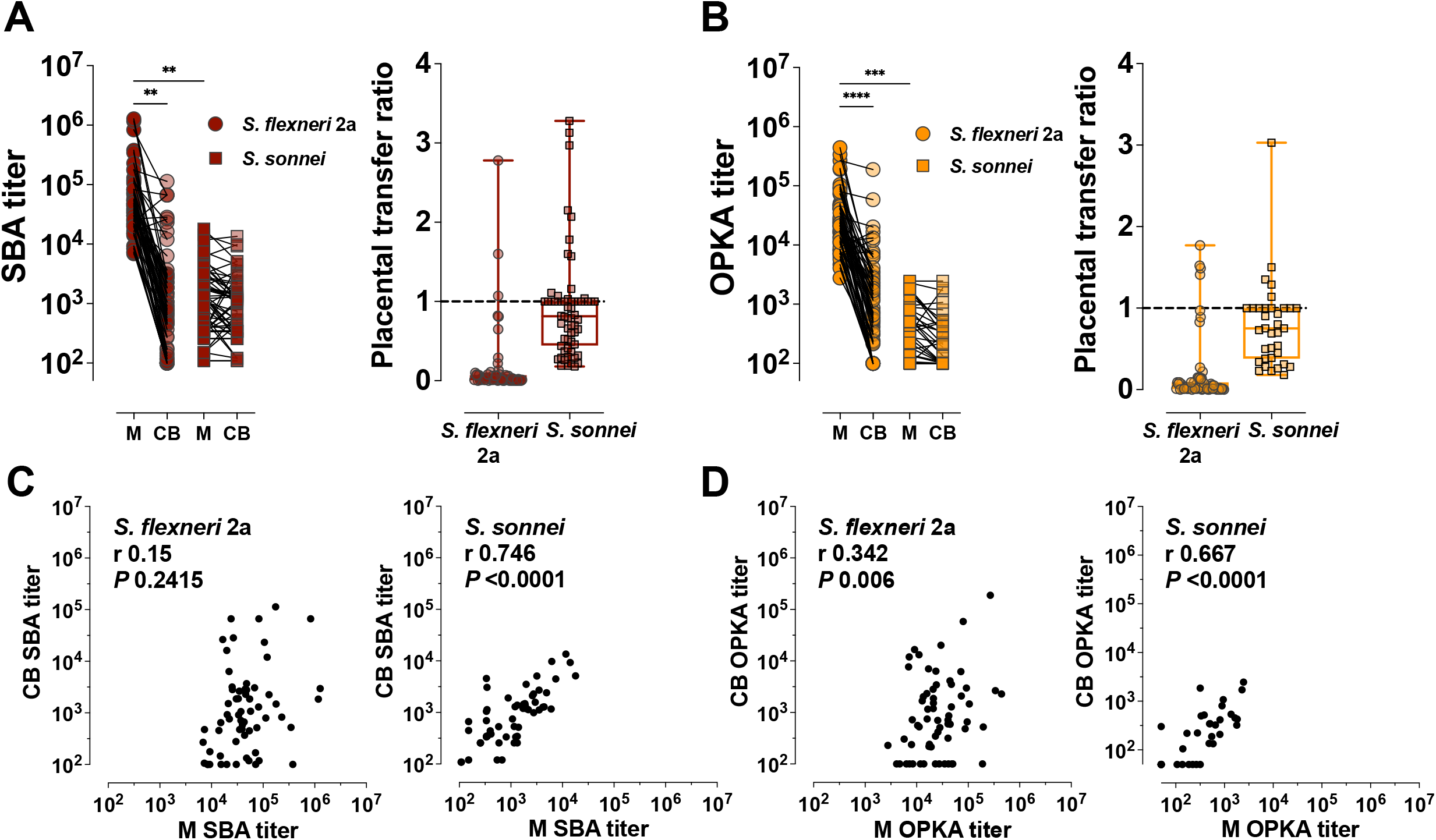
Maternal and placentally-acquired infant functional antibodies against *Shigella*. (A) Serum bactericidal antibody (SBA) and (B) opsonophagocytic killing antibody (OPKA) titers measured in maternal (M) and cord blood (CB) sera (left graphs). Data represent individual titers. Placental transfer ratios (cord blood titer/maternal titer) for each dyad are shown on the right. Asterisks indicate statistically significant differences as determined by one-way ANOVA with a Tukey’s post-test correction; ** *P* < 0.01, *** *P* < 0.001, **** *P* < 0.0001. Whiskers indicate minimum and maximum values. Line is at ratio = 1. (C) and (D) Associations between maternal (M) and cord blood (CB) SBA and OPKA titers, respectively. Pearson’s r and *P* values are shown within each graph.

### Specificity of maternally acquired functional antibodies

Mouse monoclonal antibodies specific for *Shigella* LPS were reported to have bactericidal activity (27). Several studies have reported increases in SBA titers in response to *Shigella* polysaccharide-based vaccine candidates in adult volunteers (28-30). LPS is therefore presumed to be the main antigenic target of antibody-mediated shigellacidal activity. It is not known, however, whether antibodies with other specificities deploy or contribute to this antimicrobial function. To address this question, we probed the antigen-specificity of the SBA activity in our maternal and cord blood sera by evaluating complement-dependent *Shigella* killing in samples that had been depleted of specific antibodies by pre-adsorption with increasing amounts of *Shigella* antigens; the efficiency of antibody removal was demonstrated by a decrease of ELISA binding signal of the adsorbed sera (Supplementary Figure 1A and B). Depletion of *S. flexneri* 2a LPS antibodies from both maternal and cord blood serum resulted in a proportional (dose-responsive) reduction of bactericidal activity that was serotype-specific i.e., adsorption of *S. sonnei* LPS antibodies did not reduce *S. flexneri* 2a LPS killing (Figure 5A and B). Removal of IpaB-, IpaC-, IpaD-, IpaH- and VirG-specific antibodies had no effect on *S. flexneri* 2a complement-mediated killing. To further ascertain the contribution of these antigenic targets in antibody-mediated bactericidal activity, we examined the capacity of antibodies in maternal and infant sera to kill *S. flexneri* 2a BS103, an isogenic strain lacking the invasion plasmid (31) (and which therefore does not express Ipa proteins and VirG) alongside WT *S. flexneri* 2a. Bactericidal killing curves and SBA titers were similar regardless of target strain (Figure 5C and D). These results suggest that LPS is the primary target for antibody-mediated bactericidal activity.

**Figure 5.**
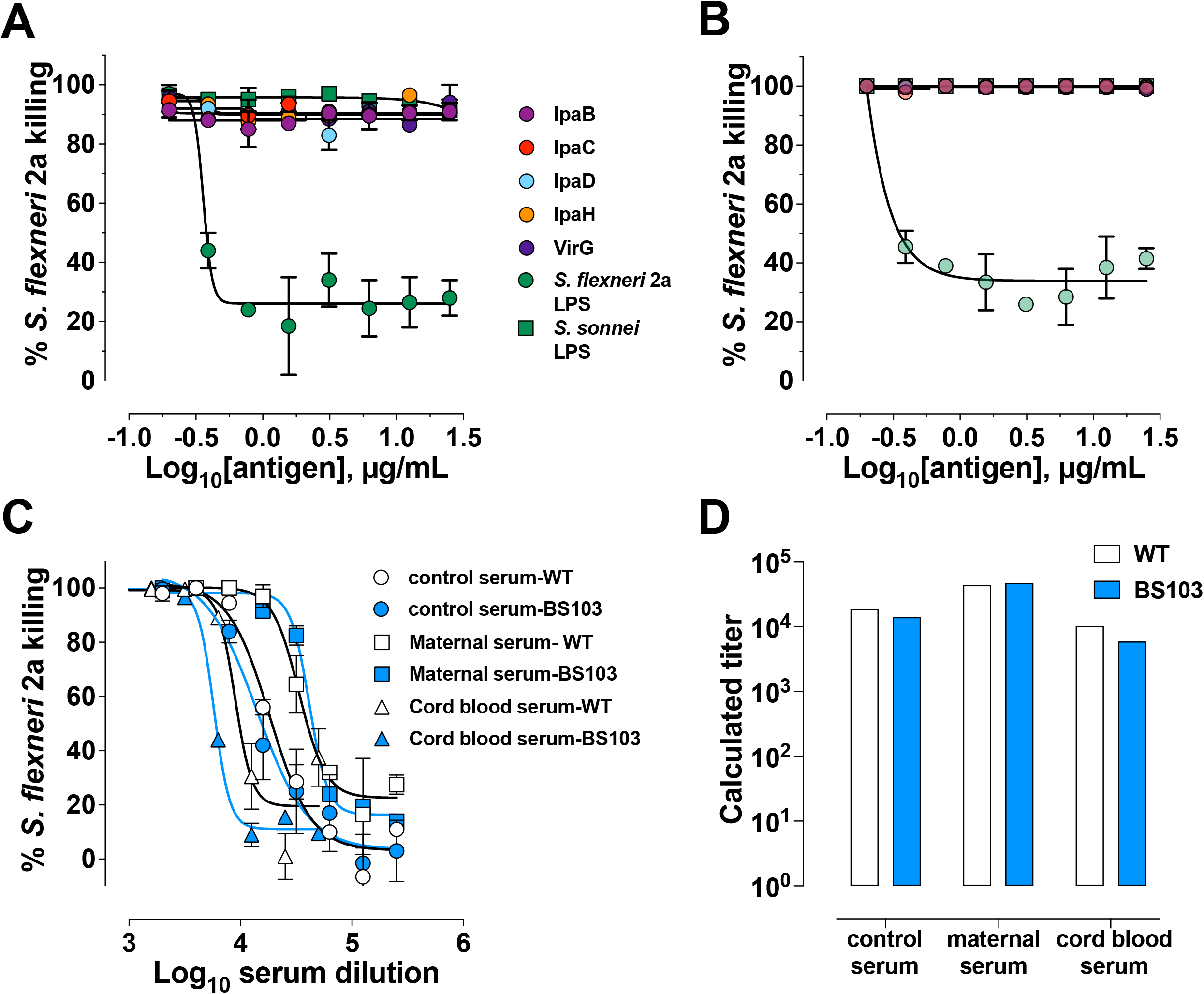
*Shigella* LPS-specific antibodies are the main contributors of bactericidal activity. Percent killing of *S. flexneri* 2a in a representative maternal (A) and cord blood (B) serum pair depleted of antigen-specific antibodies. (C) Killing curves of either wild-type (WT) *S. flexneri* 2a 2457T (clear symbols) or *S. flexneri* 2a BS103 (virulence plasmid-cured strain, shaded symbols) by hyperimmune control serum (control) and a representative maternal or cord blood serum pair at different dilutions. (D) Bactericidal titers of serum against *S. flexneri* 2a WT (clear bars) or BS103 (shaded bars).

### Antibody isotypes that mediate bactericidal activity in mothers and infants

Having identified LPS as the molecular target of antibody function, we compared LPS IgG and SBA titers in both maternal and infant serum. A positive and significant correlation was observed between maternal *S. sonnei* LPS IgG and SBA (Pearson’s r = 0.686) but the same was not true for *S. flexneri* 2a (Pearson’s r = 0.281) (Figure 6A). However, in the infants, LPS IgG and SBA titers were positively and significantly correlated for both *S. flexneri* 2a (Pearson’s r = 0.614) and *S. sonnei* (Pearson’s r = 0.707) (Figure 6E). These results hinted that another component, other than IgG, was contributing to maternal *S. flexneri* 2a SBA but was not sieved through the placenta.

**Figure 6.**
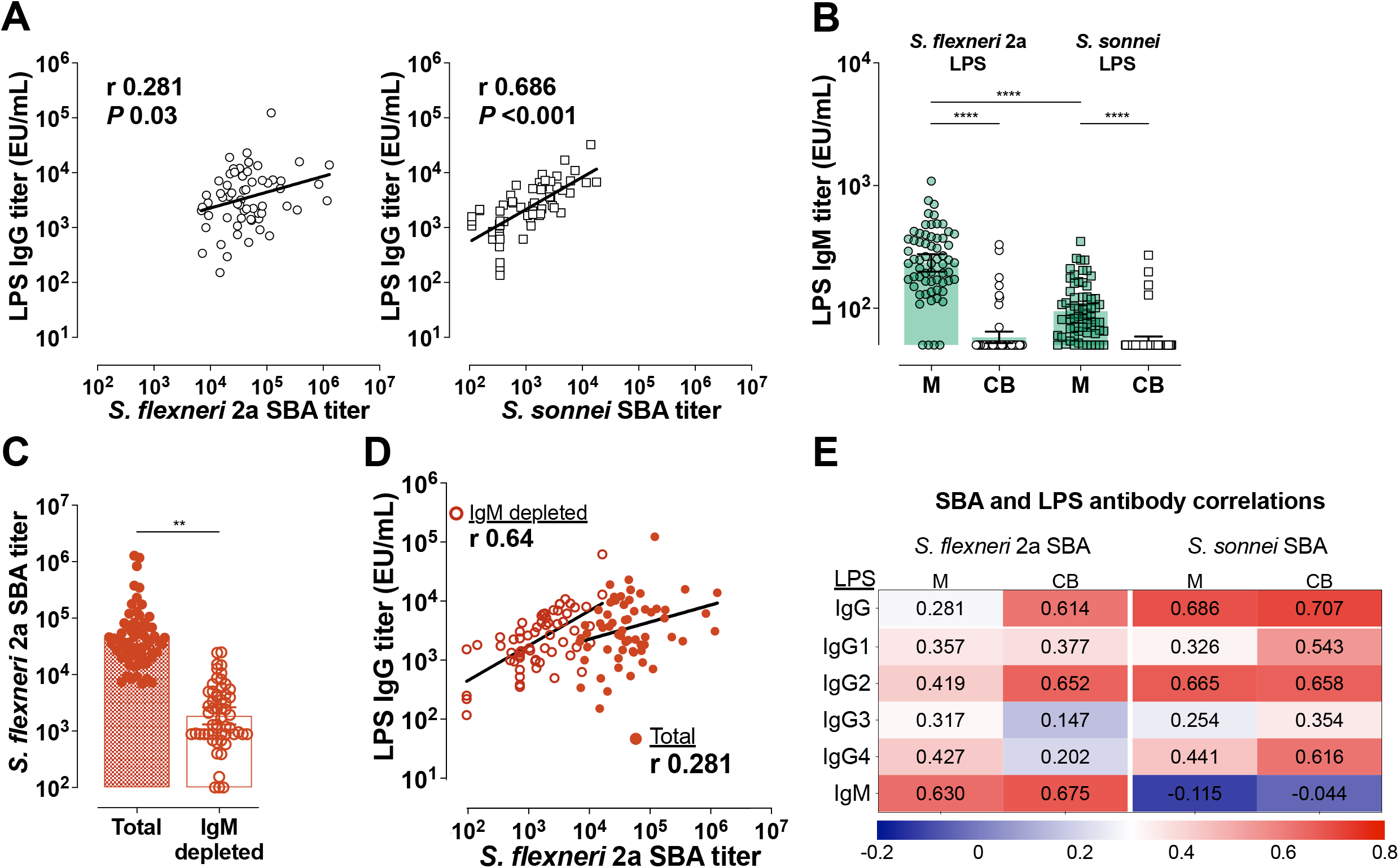
*Shigella* LPS-specific IgG and IgM mediate bactericidal activity. (A) Correlations between maternal LPS IgG and SBA for *S. flexneri* 2a (left panel) or *S. sonnei* (right panel). (B) Mean IgM titers (bars) against *S. flexneri* 2a and *S. sonnei* LPS in maternal (M) and cord blood (CB) sera. Symbols represent individual titers. Maternal and cord blood titers were compared by one-way ANOVA with a Tukey’s post-test correction; **** *P* < 0.0001. (C) Mean SBA titers (bars) in maternal serum before (Total) or after IgM depletion (IgM depleted). SBA titers in the two groups were compared by paired t-test; ** *P* < 0.01. Symbols represent individual titers. (D) Associations between *S. flexneri* 2a LPS IgG and SBA titers before (filled circle) and after IgM depletion (open circles). Pearson’s r values are shown within the graph. (E) Heatmap showing associations (Pearson’s r) between SBA and Total IgG, IgG subclasses and IgM against LPS.

IgM is a strong activator of complement that could account for the excess maternal *S. flexneri* 2a SBA and OPKA observed. LPS-specific IgM titers against both *S. flexneri* 2a and *S. sonnei* LPS were detected in maternal serum and in a handful of cord blood samples by ELISA (Figure 6B). While similar in the infants, there was substantially higher IgM against *S. flexneri* 2a as compared to *S. sonnei* LPS (mean of 263 EU/mL compared to 99 EU/mL) in maternal blood (Figure 6B). Depletion of maternal IgM greatly diminished *S. flexneri* 2a SBA (Figure 6C). *S. flexneri* 2a SBA titers measured in the IgM-depleted maternal sera were strongly associated with maternal LPS-specific IgG (Pearson’s r = 0.64); this was in contrast to the limited correlation observed when SBA was measured in intact (IgM- and IgG-containing) sera (Figure 6D). These results attribute complement dependent *Shigella* killing activity to both LPS-specific circulating IgG and IgM. In the correlation analyses, maternal *S. flexneri* 2a SBA was more closely associated with LPS IgM (likely due to its abundance in sera) while *S. sonnei* SBA was mostly associated with LPS IgG (Figure 6E). Associations were also calculated for SBA and LPS-specific IgG, IgG1-4, and IgM for both strains in maternal and cord blood. IgG2 (the predominant LPS-specific antibody) was the subclass most associated with *S. sonnei* and *S. flexneri* 2a SBA in cord blood serum (Figure 6E). IgG1 against *S. sonnei* LPS was equally associated with SBA in the infants.

### Comparisons with protective titers

Finally, to place the mother-infant antigen-specific and functional antibody titers determined in this study in the context of protective immunity, we compared serological outcomes in the dyad with those measured in individuals who remained healthy or had only mild disease when challenged with wild-type *S. flexneri* 2a in a CHIM study (11). IpaB and *S. flexneri* 2a LPS-IgG titers in maternal serum and cord blood were significantly higher than those found in adult American volunteers that remained healthy post experimental oral challenge (Figure 7A). Likewise, maternal *S. flexneri* SBA and OPKA titers were similar or higher than those of the same protected individuals. In contrast, the functional SBA and OPKA antibodies in the infants were significantly lower than those of volunteers clinically protected against experimental *Shigella* infection (Figure 7B).

**Figure 7.**
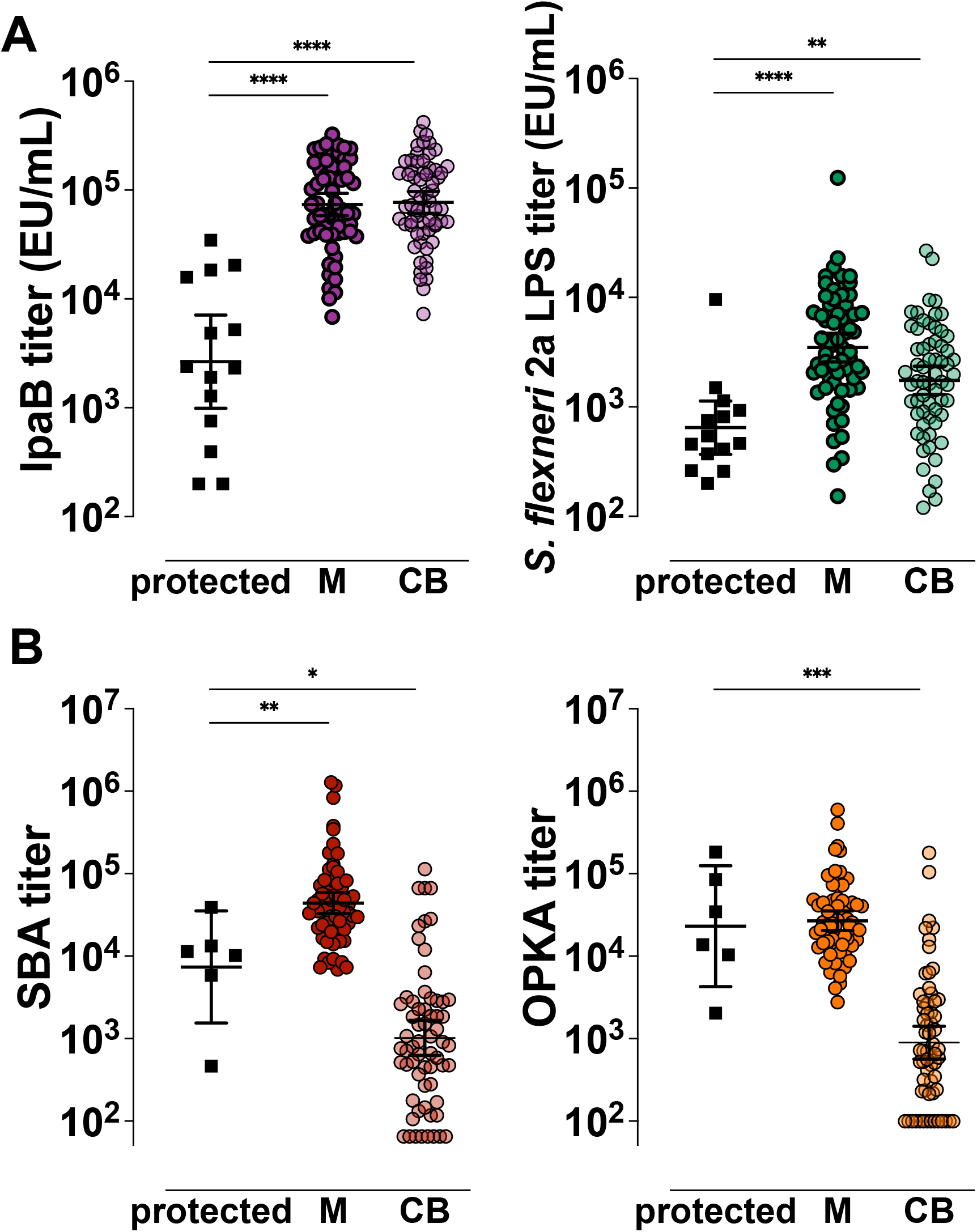
Comparative analysis of *Shigella* antigen-specific and functional antibody titers in the mother-infant dyad in relation to clinical protective thresholds. IpaB- and *S. flexneri* 2a LPS-specific (A) and *S. flexneri* 2a functional (B) serum antibody titers measured pre-challenge in North American volunteers (black squares, protected) that remained healthy or had mild disease after wild-type *S. flexneri* 2a infection and in Malawi maternal (M) or cord blood (CB) sera. Symbols represent individual titers. Differences between groups were determined by paired t-test; *P* > 0.05, * *P* < 0.05, ** *P* < 0.01, *** *P* < 0.001, **** *P* < 0.0001.

## Discussion

Through lifelong exposure, adults living in endemic regions develop natural immunity against *Shigella*, which has been attributed to antibodies against serotype specific LPS (10). The incidence of moderate to severe diarrhea attributable to *Shigella* progressively increases after the first year of life, reaching its peak in young children 24 to 59 months of age. Although they experience other diarrheal diseases, young infants are shielded from *Shigella* dysentery presumably through maternal immunity acquired via placenta or breast milk (16). The exact elements that prevent infection in these children, the contribution of antibodies specific for LPS or for other bacterial antigens, and the antimicrobial mechanisms involved are not known. An understanding of the particular components of this shielding immunity is important as new candidate vaccines targeted for the susceptible infant and toddler groups continue to advance in the clinical pathway. To identify maternally derived humoral immune components that may contribute to protection of young infants during the first months of life, we characterized the repertoire of *Shigella*-specific antibodies in a cohort of Malawian mothers and their infants at the time of birth.

Serum IgG specific for both *S. flexneri* 2a LPS and *S. sonnei* LPS were detected in maternal and in cord blood sera, although their placental transfer efficiency was moderate, 0.51 and 0.57, respectively. Two previous studies of transplacental transfer of antibodies against *S. flexneri* 2a and *S. sonnei* LPS in Israel (32) and against *S. sonnei* LPS in Vietnam (21) also found high levels of IgG specific for *Shigella* LPS in maternal sera. Our data suggest that the mothers in our cohort had been exposed to both *S. flexneri* 2a and *S. sonnei*, which is consistent with the reported serotype prevalence in countries neighboring Malawi (20, 33), and other endemic regions (32, 34, 35). The study in Vietnam reported a much higher median transfer ratio (1.33) for *S. sonnei* LPS IgG (21) than was observed in our cohort, which may reflect regional differences in seroprevalence due to *Shigella* circulation (and possibly increased proportions of LPS-specific IgG1).

An abundance of serum IgG against *Shigella* type 3 secretion system (TTSS) proteins IpaB, IpaC, IpaD, IpaH, and the virulence factor VirG were observed in maternal and infant circulation. This is a novel finding; such a detailed serological interrogation of protein-specific antibodies in field or clinical studies had been hindered by the difficulty in obtaining *Shigella* antigens of high quality and in sufficient yield; classical literature that report Ipa antibody analyses in endemic regions typically relied on crude and undefined protein extracts (9, 12). Different from antibodies to LPS, protein-specific antibodies were efficiently transferred to the newborns; cord blood IgG titers were similar or even higher than those in maternal sera for all the proteins examined. Consistent with our findings, a reduced placental transfer of polysaccharide-compared to protein-specific antibodies has been reported for other pathogens, such as *Haemophilus influenzae, Neisseria meningitidis*, and *Streptococcus pneumoniae* (36-38). Despite the differences in magnitude, we observed that maternal and infant antibody levels were correlated for all antigens, which confirms the regulated and selective nature of transplacental transfer in a process that is antigen/antibody dependent. *Shigella*-specific placental antibody sieving was distinctly linked to IgG subclass. While protein-specific antibodies were primarily IgG1 (followed by IgG2, IgG3, and IgG4), *S. flexneri* 2a and *S. sonnei* LPS-specific IgG contained mainly IgG2 (followed by IgG1, IgG3, and IgG4). The superior levels of protein-specific IgG1 in infant blood as compared to LPS-specific IgG2 is consistent with the hierarchy of receptor-mediated IgG subclass transport based on affinity to FcRn and other placental Fc receptors (37-39). Given their distinct functional attributes, the IgG subclass profile available to the infants will determine the antimicrobial capacity of humoral immunity early in life (1, 23, 43).

SBA and OPKA titers have been correlated with clinical protection in adults experimentally infected with virulent *S. flexneri* 2a (11). Our evaluation of bactericidal activity specificity—both the antibody depletion experiments and SBA using the plasmid-cured isogenic S. *flexneri* 2a BS103—revealed the preeminence of LPS antibodies in antibody-mediated complement dependent bacterial killing in maternal and infant sera. Consistently, LPS IgG (and particularly IgG2) was strongly correlated with SBA activity for both *S. flexneri* 2a and *S. sonnei*. Interestingly, IgG2, which makes up the bulk of LPS-IgG, is a poor complement activator. However, IgG2 is known to mediate complement-dependent killing of *Haemophilus influenzae* type b, albeit not as efficiently as IgG1 (40). On the other hand, LPS IgG2 may activate an alternate pathway when epitope densities are high (41), which is likely the case for a surface-exposed target like LPS. IgG2 can act in a complement-independent manner as has been shown with opsonophagocytic activity against *S. pneumoniae* (42), which is consistent with OPKA activity observed in our study. Antigenic targets of antibody-mediated bactericidal killing other than LPS as well as other protein-specific antibody-dependent functions need further investigation.

Unexpectedly enhanced SBA and OPKA activity against *S. flexneri* 2a was observed in the mothers (but not in the infants) from our cohort that was linked to high levels of IgM (a potent activator of complement that is not placentally transferred). A similar reduction in functional activity in infant compared to maternal sera, also due to maternal IgM, has been shown against *E. coli* and *Salmonella* (43, 44). Though mothers in our cohort had comparable levels of LPS IgG against both *Shigella* serotypes, maternal LPS-specific IgM against *S. flexneri* 2a was markedly higher resulting in heightened SBA activity. Differences in IgM levels likely reflect frequency of exposures and strain circulation, suggesting, in our study, a higher prevalence of *S. flexneri* 2a as compared to *S. sonnei* in the Blantyre, Malawi, region. A handful of infant samples had detectable IgM against LPS from both serotypes. IgM against environmental and vaccine antigens has been reported in cord blood from infants in LMIC but not in those from industrialized nations. The origin of this IgM is unclear and presumed to reflect environmental factors, such as intrauterine infections that affect placenta integrity (45, 46), and non-specific natural antibodies (45, 47). Maternal SBA and OPKA against both serotypes were strongly associated implying shared antibody contribution to microbial killing.

The underlying premise for dissecting the humoral immune profile against *Shigella* in mothers and infants living in endemic regions is the epidemiological evidence of the lowest risk of infection in these groups. Maternal *S. flexneri* 2a LPS-IgG, SBA, and OPKA titers were comparable or higher than those observed in North American volunteers who remained healthy following challenge with wild-type *S. flexneri* 2a organisms (11). The functional antibody activity in the infants was noticeably below the threshold of clinically protected adults. In contrast, serum IgG against protective target antigens IpaB and VirG in mother and infant sera were well above those detected in the clinically protected challenged volunteers (11). The low levels of LPS-specific IgG and their limited functional capacity (SBA and OPKA) in cord blood, along with high levels of maternal protein-specific IgG and its efficient transfer, argue in favor of a more prominent role of antibodies against *Shigella* virulence antigens in preventing *Shigella* infection than originally thought. Similarly, others observed that naturally acquired maternal antibodies against pneumococcal proteins, unlike anti-polysaccharide antibodies, were associated with protection against nasal carriage in infants during the first 3 months of life (48). Further studies are warranted to dissect the mechanisms, not captured by our traditional assays, by which these protein-specific antibodies block microbial infection and to corroborate their disease protective capacity in humans.

A *Shigella* vaccine that is efficacious in children under 3 years of age, the most vulnerable target group, would make a major public health impact. The limited efficacy of a clinically advanced O-polysaccharide-based vaccine candidate for young children (49, 50) has been linked with young children’s hypo-responsiveness to *Shigella* LPS (and likely impaired bactericidal/phagocytic antibody activity). Our results showing the abundance of protein-specific antibodies in groups naturally immune (low-risk group) to *Shigella* and the ease with which children respond to protein-based immunization highlight the prospect of a *Shigella* protein-based vaccine approach. Purified IpaB and IpaD (51-53) or a formulation containing IpaB and IpaC (Invaplex, (54)) have been shown to prevent *Shigella* infection in preclinical studies. While the exact operative mechanisms are yet to be determined, antibodies targeting *Shigella* proteins are expected to block *Shigella* attachment and TTSS protein translocation (anti-IpaB, -C, and -D), prevent bacterial spread and replication (anti-VirG and anti-IpaH), and prevent inflammation (55). A vaccine combining multiple conserved *Shigella* proteins would not only be likely broadly protective, but also effective, targeting multiple mechanisms important for *Shigella* invasion and virulence.

One limitation in our study was that maternal serum was obtained within 3 months of birth. While some antibody features may change after birth, we did not find significant differences in maternal titers determined in sera collected prior to or after delivery. The limited sample size may have also precluded more extensive and complex statistical analyses. It would be important to conduct similar analyses of the antibody repertoire beyond birth and through the first 3 years of life to better understand trends of disease and immune acquisition, and to investigate age-specific antibody mediated antimicrobial functions using age-relevant immune cells (17) to recreate protective elements that would operate in vivo.

In summary, we have demonstrated, for the first time, the efficient placental transfer of maternal antibodies against *Shigella* protein antigens and their availability at high levels to the infant at birth along with the less efficient transfer of LPS-specific IgG with bactericidal and opsonophagocytic killing activity. Our results define the maternally acquired protective antibody repertoire available to infants at birth and suggest a larger role for protein-specific immunity than has previously been appreciated. Exploring the concept of a protein-based vaccine that would target these antigens either alone or in conjunction with *Shigella* LPS is warranted. Finally, our findings also emphasize the need to better understand immunity in young children to inform preventive strategies. An in-depth interrogation of adaptive immunity accrued from infection during early childhood as well as offered through breastmilk would help identify protective elements in this most vulnerable group.

## Acknowledgments

We are grateful to the women who volunteered to participate and to the nurse midwives of the Mfera Health Centre maternity ward and antenatal clinic who supported this study. We also thank Dr. Edwin Oaks for critical reading of the manuscript and advice on Ipa experiments. We thank Syze Gama and Dr. Avital Shimanovich for providing technical and logistic expertise during the sample collection process and members of the Pasetti Lab for technical assistance and discussion.

## Methods

### Study population and sample collection

Our study population consisted of mothers and infants recruited between January and November 2016 at Mfera Health Clinic in Chikwawa, Malawi. Healthy pregnant women who were HIV seronegative were enrolled either at the antenatal clinic or during delivery. Infants were enrolled at birth. Baseline information on mother and the infant’s health was obtained at enrollment. Other information such as village, baseline health info, and current physical complaints was also collected during the visit. Umbilical cord blood was collected at delivery at the Mfera Health Clinic. Venous maternal blood was obtained at enrollment, either during screening before birth, at a well-child visit (week 1, 6, or 10) or at 3 months. Samples were frozen and shipped to the University of Maryland School of Medicine in Baltimore for analysis. This study was approved by the Institutional Review Board of University of Maryland School of Medicine, and the College of Medicine Research and Ethics Committee (COMREC) at the College of Medicine in Malawi. All participating mothers provided written informed consent for themselves and their infants. Serum samples from North American individuals were obtained from a previous clinical study performed at the Center for Vaccine Development (University of Maryland, Baltimore) under approved IRB protocols. Serum samples tested were obtained at day -1, prior to challenge with 1×10^3^ CFU of the wild-type *S. flexneri* 2a strain 2457T as described previously (56); some of these volunteers had been previously vaccinated and had varying degrees of immunity. Specimens were selected from volunteers who remained healthy or who experienced mild disease, as previously described (11).

### Antigen-specific antibody analysis

*Shigella* antigens IpaB, IpaC, IpaD, *S. flexneri* 2a LPS and *S. sonnei* LPS were obtained from Walter Reed Army Institute of Research (WRAIR). The N-terminal domain of VirG was expressed and purified inhouse in an *E. coli* expression system (Chitra STS et al., unpublished). Immune responses to IpaH were measured using the conserved C-terminal domain of IpaH1.4 (IpaH-CTD) produced by Vaxcyte (N. Kapoor et al., manuscript submitted for publication). IpaH1.4 was selected because it was one of the top isoforms recognized by serum from vaccinated or *S. flexneri* 2a-challenged individuals using a core *Shigella* proteome microarray (57). Antigen-specific serum IgG titers were measured by ELISA as previously described (58). Briefly, Immulon 2HB plates (Thermo Scientific, Waltham MA) were coated with IpaB, IpaC, IpaD, and IpaH at 0.1µg/mL in PBS, and VirG, *S. flexneri* 2a LPS and *S. sonnei* LPS at 5µg/mL in carbonate buffer, pH 9.6. Plates were incubated for 3h at 37°C and blocked at 4°C overnight in PBS containing 10% w/v non-fat dry milk (NFDM). Sera diluted in PBS containing 10% NFDM and 0.05% Tween-20 (PBS-T) were added, and the plates incubated at 37°C for 1h. Plates were incubated with HRP-labeled goat IgG specific for human IgG (Jackson Immuno Research, West Grove, PA) for another 1h at 37°C. Plates were washed 6 times with PBS-T following every incubation step. Tetramethylbenzidine (TMB; KPL, Gaithersburg, MD) was added as substrate for 15 min in the dark with shaking, and the reaction was stopped by adding 1M phosphoric acid (Millipore Sigma, Burlington, MA). Endpoint titers were calculated as the inverse serum dilution that resulted in an absorbance value at 450 nm of 0.2 above background and were reported as the ELISA units/mL.

Antigen-specific IgG subclasses were measured using *Shigella* multiplex assay using the MesoScale Diagnostic platform (MSD, Rockville, MD). Assays were run in the same way as the antigen-specific ELISAs, with a few exceptions: 1. There was no antigen-coating step as the antigens were pre-printed on MSD plates. 2. After the serum incubation step, the plates were incubated with biotinylated anti-IgG subclass antibodies (SouthernBiotech, Birmingham, AL) plus the SULFOTag-STREP (MSD) for another 1h at 37°C. 3. Binding was then detected using MSD GOLD Read buffer. The plates were read using an MSD sector imager, model 2400 and data analyzed by the MSD workbench software provided by the manufacturer. The Electro Chemiluminescence signal (minus background from blank) for each sample (diluted at 1:100 in PBS containing 10% NFDM) was reported.

### Antibody functional analysis

#### Bacterial growth conditions

For both SBA and OPKA, *S. flexneri* 2a 2457T and *S. sonnei* 53G were grown as previously described (11). Briefly, bacteria were streaked onto Tryptic Soy agar (TSA) plates supplemented with Congo Red and incubated overnight at 37ºC. Single red colonies were propagated in LB media and grown to early log phase, conditions that support the growth of stable virulent organisms. *S. flexneri* 2a strain BS103 (avirulent plasmid-cured strain) was a gift from Dr. Robert Kaminski (Walter Reed Army Research Institute); bacterial cultures were propagated to early log phase from single white colonies picked from TSA plates supplemented with Congo Red.

#### SBA

The SBA assay was performed as previously described (59). SBA titers were determined by Opsotiter (59) as the reciprocal of the serum dilution that produced 50% bacterial killing as determined by Reed-Muench regression analysis. The provisional reference serum sample, Korean QC19, was run with each assay to normalize and reduce variability between assays. Korean QC19 was assigned a titer = 28000 for *S. flexneri* 2a and 1100 for *S sonnei* (59). The lowest dilution tested was 1:200 so that the lowest titer is 100. For *S. flexneri* 2a BS103, SBA assays were performed in a similar manner, except that LB agar plates spotted with 10 µL of the reactions were grown overnight at 37ºC.

#### OPKA

OPKA was also performed as previously described (11) with some modifications. Briefly, 10 µL of target bacteria (∼10^4^ CFU/mL) was opsonized by mixing with 20µL of a heat-inactivated, serially diluted test sample in a well of a round-bottom microtiter plate and incubated for 15 min at 37°C in room air, with shaking. As with the SBA, control wells had bacteria, baby rabbit complement (BRC), and buffer only; no test sample was added to these wells. 10µL of baby rabbit complement (10% final concentration), mixed with 60µL of 10^5^ dimethylformamide; (DMF)-differentiated HL-60 cells (ATCC CCL-240) were then added to the reaction mixture (for a 100µL total volume). Following a 45 min incubation at 37°C, 5% CO2, 10 µL from each well was spotted on LB agar. The agar plates were incubated overnight at 29°C for *S. flexneri* 2a and 26°C for *S. sonnei*. The percentage of bacteria that were phagocytosed and killed per well was determined by measuring the colony counts and determining the titer, as was done for the SBA (above). The Korean QC19 provisional reference sample was included in each assay and found to have an average titer of 15574 for *S. flexneri* 2a and 288 for *S sonnei*. The lowest dilution tested was 1:200; samples with titer below 200 were assigned an arbitrary value of 100.

#### Antibody depletion

(i) LPS and protein-specific antibodies were removed by incubating serum samples on ELISA plates coated with increasing antigen concentrations (0 - 25µg/mL) for 1h at 37°C with shaking. Reduced antibody content in adsorbed samples was confirmed by ELISA. The antibody-depleted sera were then tested for SBA activity as described above. (ii) IgM was removed by incubating serum samples with 50mM of beta-mercaptoethanol for 1h at 37°C. The IgM-depleted serum was further diluted and used in the SBA reaction as described above.

### Statistical analysis

Geometric mean titers (GMT) were calculated for antigen-specific IgG in maternal and cord blood sera. Comparisons of titers between maternal and cord samples, or between antibody levels against different antigens, were measured by paired t-test or one-way analysis of variance (ANOVA) with Tukey’s post-test correction. Placental transfer ratios were assessed as a ratio of cord blood divided by maternal antibody titers. Associations between maternal and cord blood titers were calculated using Pearson’s correlation. A two-way ANOVA with Tukey’s post-test correction was used to compare transfer ratios between IgG subclasses. All statistical analysis was conducted using GraphPad Prism 9.

