## Supplementary figures and images for "Repertoire of naturally acquired maternal antibodies transferred to infants for protection against shigellosis"

### Supplementary Figure 1

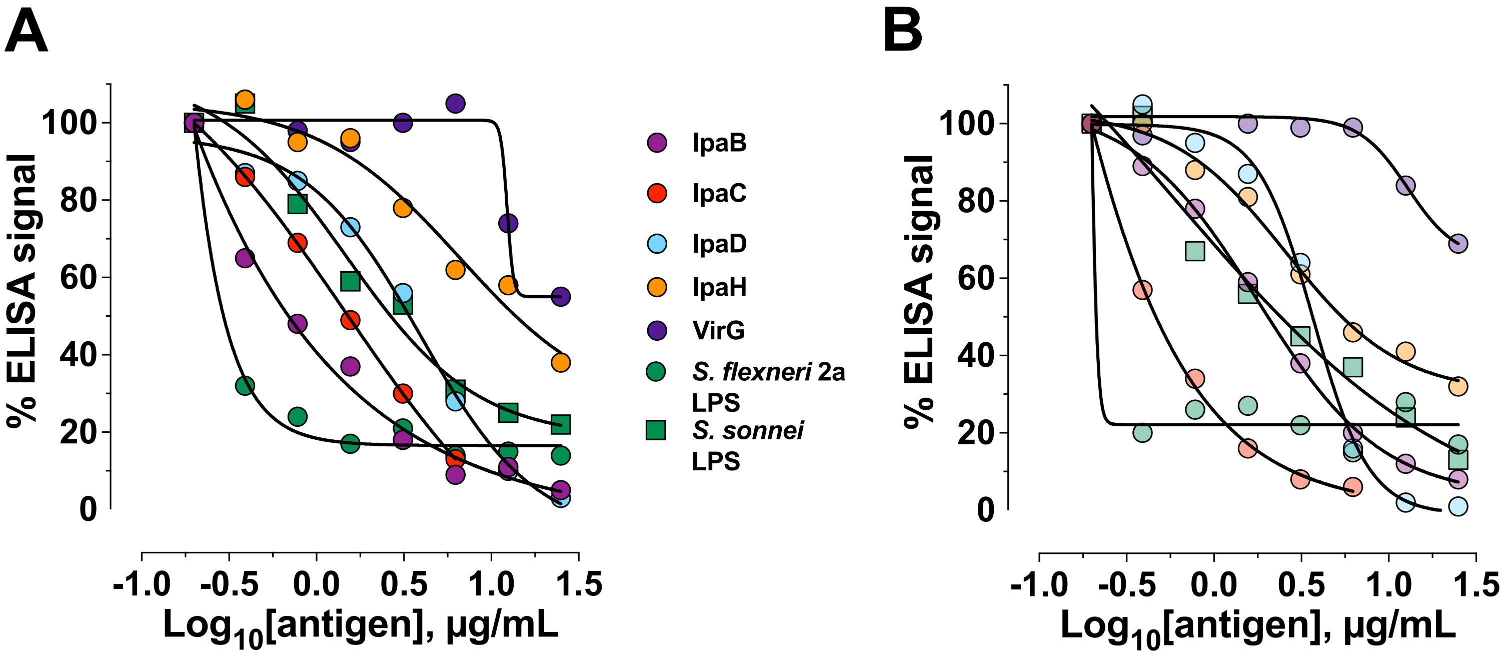

### Supplementary Figure 2

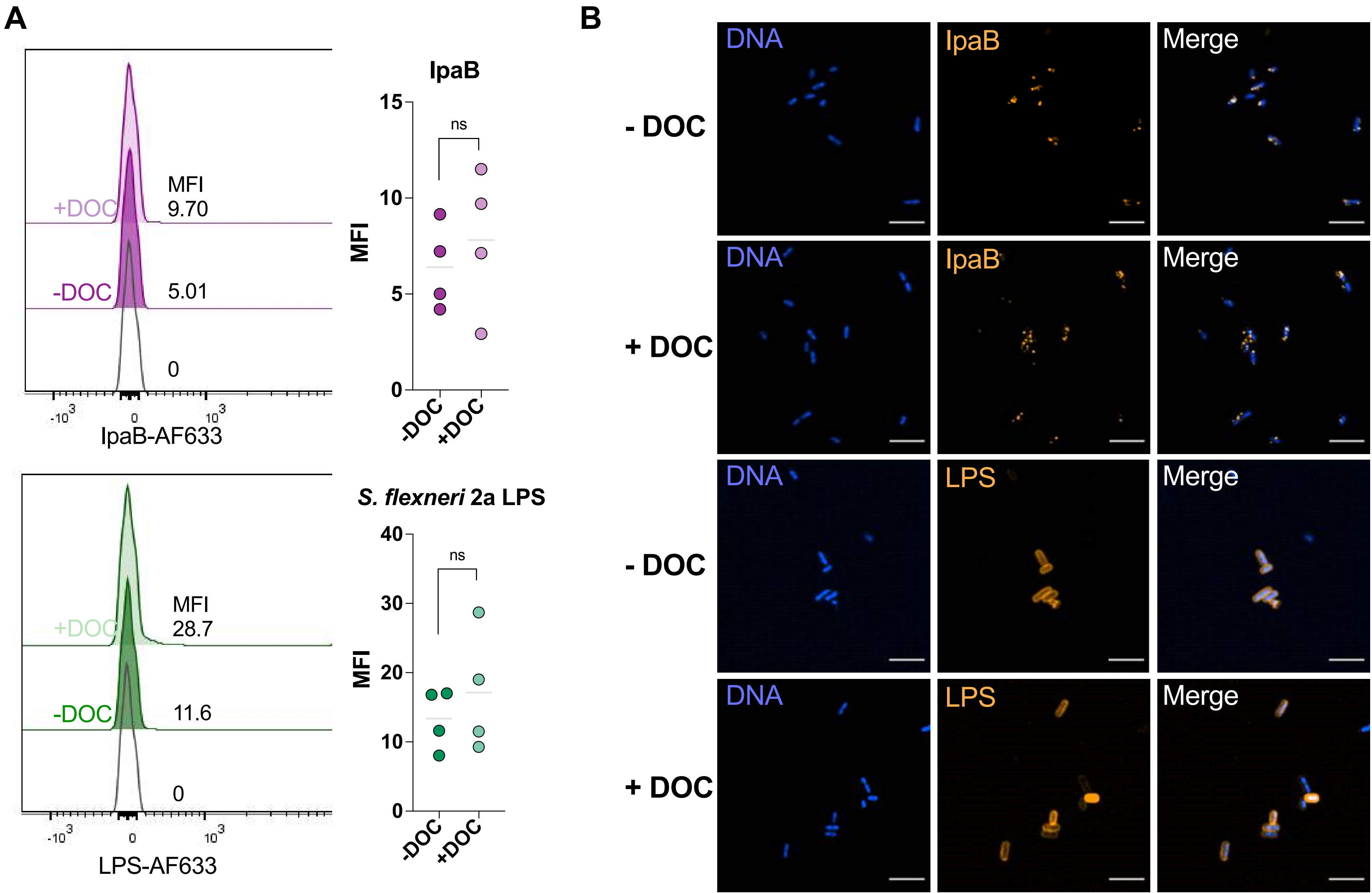
