## Supplementary materials for "Repertoire of naturally acquired maternal antibodies transferred to infants for protection against shigellosis"

**Supplementary Tables**

| **Supplementary table 1. Spearman correlation between Log_10_-Transformed Anti-*Shigella* antigen–specific IgG Transfer ratios and maternal and infant covariates** | | | | | | | | | | |
| --- | --- | --- | --- | --- | --- | --- | --- | --- | --- | --- |
|  | Maternal serum (Log_10_ titer) | | Maternal age | | Parity | | Gestational age | | Infant birthweight | |
|  | r | *P* | r | *P* | r | *P* | r | *P* | r | *P* |
| *S. flexneri* 2a LPS | -0.25 | **0.05** | 0.05 | 0.68 | -0.20 | 0.15 | -0.10 | 0.43 | 0.02 | 0.87 |
| *S. sonnei* LPS | -0.16 | 0.22 | -0.06 | 0.63 | -0.21 | 0.13 | -0.13 | 0.30 | 0.18 | 0.15 |
| IpaB | -0.26 | **0.04** | -0.01 | 0.96 | -0.07 | 0.65 | -0.14 | 0.26 | -0.14 | 0.26 |
| IpaC | -0.27 | **0.03** | -0.03 | 0.80 | 0.11 | 0.42 | -0.12 | 0.33 | 0.13 | 0.31 |
| IpaD | 0.001 | 0.99 | -0.09 | 0.50 | 0.04 | 0.76 | -0.22 | 0.08 | -0.28 | **0.02** |
| IpaH | -0.17 | 0.18 | -0.02 | 0.88 | -0.10 | 0.47 | -0.08 | 0.55 | 0.09 | 0.49 |
| VirG | -0.38 | **0.002** | -0.08 | 0.56 | -0.17 | 0.22 | -0.09 | 0.47 | -0.07 | 0.58 |

| **Supplementary table 2. *Shigella* antigen-specific IgG subclass titers** | | | | |
| --- | --- | --- | --- | --- |
| **Antigen** |  | **Cord blood**  **Geometric Mean**  **ECL signal (range)** | **Maternal serum**  **Geometric Mean**  **ECL signal (range)** | **Transfer ratio**  **Median (range)** |
| *S. flexneri* 2a LPS | IgG1 | 243 (21-11,645) | 185 (1-2600) | 1.13 (0.57-21.04) |
|  | IgG2 | 1,632 (1-210,565) | 5,671 (5-840,470) | 0.33 (0.00-27.65) |
|  | IgG3 | 18 (1-2,466) | 25 (1-1,273) | 0.68 (0.01-1959.50) |
|  | IgG4 | 3 (1-95) | 5 (1-312) | 1.00 (0.00-85.94) |
| *S. sonnei* LPS | IgG1 | 523 (29-6,004) | 379 (15-3,002) | 1.28 (0.25-5.54) |
|  | IgG2 | 15,802 (1-287,974) | 24,142 (1-323,914) | 0.61 (0.03-16.88) |
|  | IgG3 | 19 (1-1,034) | 36 (1-1,801) | 0.53 (0.01-22.89) |
|  | IgG4 | 7 (1-405) | 16 (1-480) | 0.68 (0.01-14.73) |
| IpaB | IgG1 | 46,557 (1,869-84,196) | 45,664 (1,013-93,206) | 0.97 (0.56-1.85) |
|  | IgG2 | 7,023 (9-53,944) | 11,344 (84-109,821) | 0.53 (0.10-4.15) |
|  | IgG3 | 150 (10-6,131) | 221 (1-14,906) | 0.63 (0.01-21.55) |
|  | IgG4 | 114 (1-1,603) | 83 (1-1,651) | 1.33 (0.06-6.15) |
| IpaC | IgG1 | 30,887 (45-169,717) | 31,844 (39-227,823) | 0.99 (0.27-4.61) |
|  | IgG2 | 5,435 (1-199,595) | 8,244 (1-565,966) | 0.56 (0.06-60.83) |
|  | IgG3 | 721 (1-526,566) | 1,278 (28-741,230) | 0.60 (0.00-3.48) |
|  | IgG4 | 215 (1-4,352) | 127 (1-3,818) | 1.28 (0.01-144.96) |
| IpaD | IgG1 | 22,024 (1-481,590) | 18,317 (1-556,666) | 1.06 (0.15-56.95) |
|  | IgG2 | 1,087 (1-1,248,912) | 1,970 (1-1,096,749) | 0.68 (0.01-7.15) |
|  | IgG3 | 145 (1-5,026) | 196 (1-10,991) | 0.64 (0.01-36.00) |
|  | IgG4 | 145 (1-5,008) | 104 (1-3,398) | 1.37 (0.07-61.63) |
| IpaH | IgG1 | 98,331 (1,722-427,846) | 92,651 (942-398,831) | 1.04 (0.36-6.42) |
|  | IgG2 | 17,340 (14-856,724) | 21,868 (1-1,303,108) | 0.63 (0.05-49.28) |
|  | IgG3 | 437 (3-236,815) | 685 (17-219,837) | 0.72 (0.01-5.12) |
|  | IgG4 | 230 (1-10,748) | 230 (4-17,483) | 1.25 (0.01-16.49) |
| VirG | IgG1 | 47,804 (1,582-253,130) | 40,982 (1,250-274,979) | 1.14 (0.12-8.97) |
|  | IgG2 | 1,372 (1-143,329) | 2,156 (27-821,473) | 0.67 (0.00-878.16) |
|  | IgG3 | 603 (23-162,794) | 681 (23-323,202) | 0.89 (0.24-4.13) |
|  | IgG4 | 281 (1-17,321) | 115 (1-44,752) | 1.72 (0.14-92.20) |

ECL: Electro Chemiluminescence Signal

**Supplementary Figure Legends**

**Supplementary Figure 1. Depletion of antigen-specific antibodies in maternal and infant sera.** Percent ELISA signal (compared to non-depleted sera) in maternal (A) and cord blood (B) sera following incubation in ELISA plates coated with increasing concentration of antigen.

**Supplementary Figure 2. IpaB and LPS expression in the absence or presence of DOC.** (A) Expression of *Shigella* IpaB (top) and *S. flexneri* 2a LPS (bottom) by flow cytometry is shown in representative histograms (left) and scatter plots (right) in *S. flexneri* 2a WT grown in the absence (-DOC) or presence (+DOC) of 2.5mM sodium deoxycholate (DOC); ns, *P* >0.05. (B) Representative confocal microscopy images of expression of IpaB and *S. flexneri* 2a LPS on the bacterial surface. Scale bar = 5µm.

**Supplementary Methods**

**Bacterial growth conditions.**

*S. flexneri* 2a 2457T was streaked onto Tryptic Soy agar plates supplemented with Congo Red and incubated overnight at 37ºC. To determine whether the bile salt deoxycholate was required for expression of IpaB, single red colonies were propagated in Tryptic Soy broth (TSB) with or without 2.5mM sodium deoxycholate (DOC) and grown to early log phase.

**Flow cytometry**

Bacteria were washed twice with PBS and 50μL of equal number of cells (5x10^7^ CFU) were dispensed in separate tubes. The following mouse primary antibodies were used for staining for 1h at 4ºC, respectively: serum from mice immunized with purified IpaB (inhouse) and Hflex2a4 mAb (mouse anti-*S. flexneri* 2a LPS provided by Dr. Moon Nahm, University of Alabama). After washing, stained bacteria were incubated with goat anti-mouse Alexa Fluor Plus 633 (Thermo Fisher Scientific) diluted 1:200 in PBS for 1h at 4°C. Cells were washed and resuspended in PBS. Sample acquisition was performed on BD LSRII using FACSDiva software (BD Biosciences, USA) and antigen expression analyzed with FlowJo software (v10, Tree Star).

**Immunofluorescence staining and confocal imaging.** Bacterial suspensions (1x10^6^ CFU/ml) were placed in chamber slides (Millipore, USA) and fixed for 45 min in 4% paraformaldehyde (Electron Microscopy Sciences, USA). Bacteria were washed twice with PBS and incubated overnight at 4°C with primary mouse antibodies: serum from mice immunized with purified IpaB (inhouse) and Hflex2a4 (mouse mAb provided by Dr. Moon Nahm, University of Alabama). Antibodies were diluted 1:100 in PBS containing 5% BSA. Stained bacteria were washed twice with PBS and incubated with goat anti-mouse Alexa Fluor Plus 555 (Thermo Fisher Scientific) diluted 1:100 in PBS for 1h at RT. Slides were washed with PBS and mounted with ProLong Gold Antifade Reagent with DAPI (Cell Signaling Technology, US). Confocal imaging was carried out at the Confocal Microscopy Facility of the University of Maryland School of Medicine using a Nikon W1 spinning disk confocal microscope running NIS Elements software (Nikon). Images were captured with a 60X oil objective and settings were adjusted to optimize signal. Images were collated using FIJI/ImageJ software (NIH). Signal processing was applied equally across the entire image.
